# Finding Consensus miRNAs Silencing KLF1 Expression as A Promising Therapeutic Option of Sickle Cell Anemia

**DOI:** 10.1101/2022.11.18.517062

**Authors:** Haitham Ahmed Al-Madhagi

**Affiliations:** Biochemical Technology Program, Dhamar University, Yemen

**Keywords:** Sickle cell anemia, miRNA, gene therapy, hybridization energy, KLF1

## Abstract

Sickle cell anemia (SCA) is the almost the severest hemoglobinopathy known with no cure till date. Patients with SCA has a shorter lifespan and suffer from painful crises and end-organ damages. The goal of this in silico work is to find the consensus miRNAs targeting KLF1 gene, responsible for HbF-to-HbA switching followed by generating the corresponding miRNA sponge for gene silencing purposes. 3 publicly available databases were searched, miRDB, miRWalk and TargetScan. Afterwards, the hybridization examination of the predicted miRNAs was evaluated. Finally, the design of miRNA sponge as a means to target miRNA was performed. In conclusion, hsa-miR-330-5p was the best miRNA targeting KLF1 gene in many aspects and its miRNA sponge sequence was provided.

## 2 Introduction

Sickle cell anemia (SCA), as its name implies, a type of hemolytic anemia caused by autosomal recessive mutation in the β-chain of hemoglobin (Hb). Such simple mutation leads to huge systematic complications involving acute chest syndrome, chronic pain, recurrent infections, hemolytic anemia, nephropathy and stroke [1]. The disease shortens the lifespan of the afflicted individuals 20 years in comparison with intact people. It is estimated that approximately 3 million people worldwide suffer from SCA, with 300000 are in the US alone [2]. In addition, >1.1 million newborn babies with the HbAS (heterozygous) genotype in the US were documented [3]. The molecular basis of the disease involves the inheritance of two mutated β-chain of Hb at position 6 of the amino acid sequence, transforming the native glutamate into valine. As a consequence, the hydrophobic nature of valine attracts other hydrophobic amino acids, primarily phenylalanine, in the other subunits and, hence, clustering Hb chains together. This in turn leads to the formation of crescent or sickled shape, from which the disease gained the nomenclature (Fig 1) [4]. Indeed, the heterozygous genotype is also characterized by the precipitation of Hb molecules albeit less efficiently and prominently as the homozygous one. This takes place upon deoxygenated Hb status and restores some structure as well as function in the oxygenated form [5].

**Figure 1.**
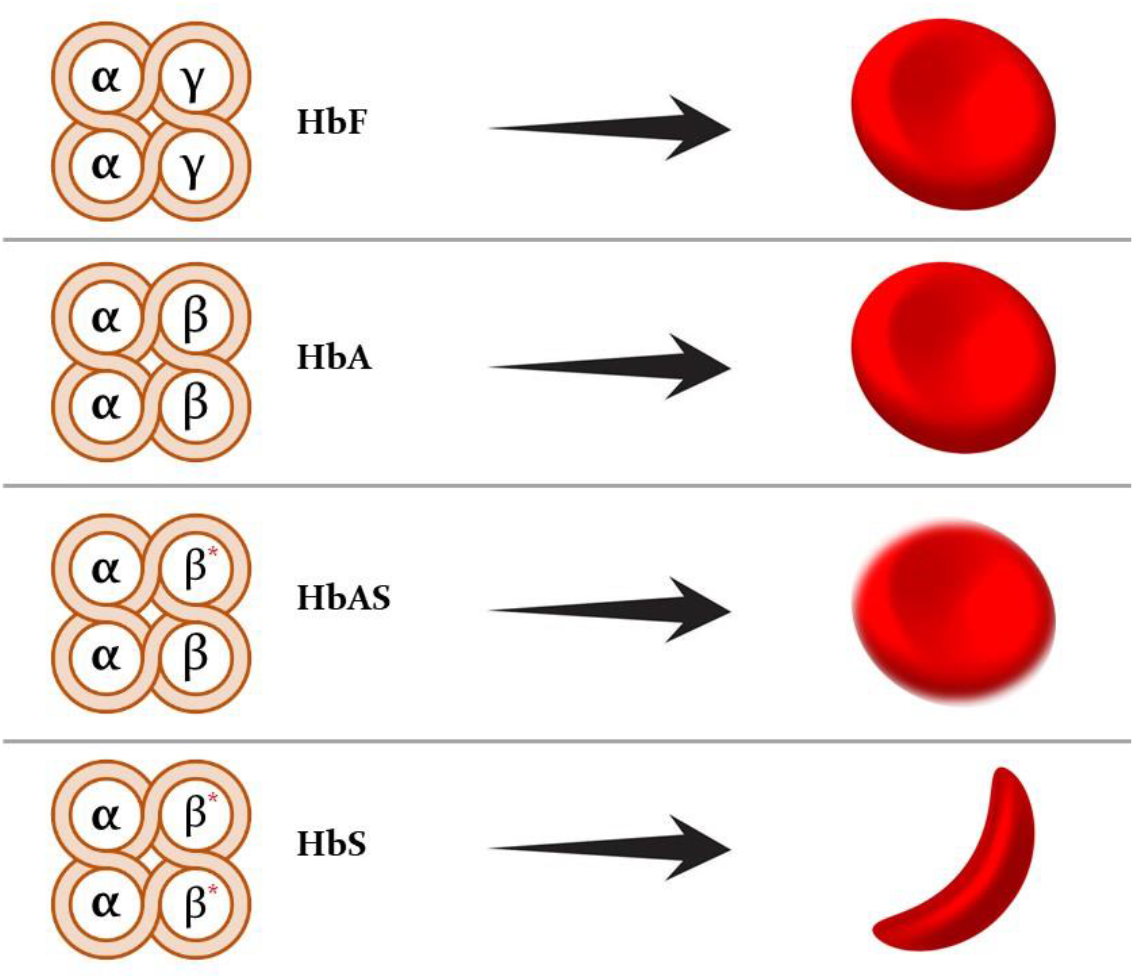
Different Hb inherited genotypes and their consequtive phenotype

The current FDA-approved first-line treatment option of SCA is hydroxyurea. Until 2017, it was the only-choice therapy of SCA. It exerts its beneficial effects mainly through upregulation of fetal Hb (HbF) which is normally functioning and can compensate for the HbS, thus restoring the oxygen-carriage function of the sickled cells. Beyond 2017, L-glutamine Oxbryta and Crizanlizumab were approved as second-option therapy demonstrating moderate positive findings [5,6]. While L-glutamine provides NH3 needed by erythrocytes to synthesize and maintain redox coenzymes (NAD and NADP) status and thereby reducing oxidative damages, Oxbryta prevents the polymerization of HbS subunits. Crizanlizumab on the other hand selectively bind P-selectin which in turn prevents the adhesion between neutrophils, platelets to endothelial cells prior occlusion of small blood vessels [7]. However, given that the disease is genetic-based, the rational and the most promising treatment is through the application of gene therapy advances. Recently, 3 main gene therapy methodologies were evaluated to test the efficitveness of the overall therapy, namely: (i) gene addition, (ii) gene editing, and (iii) gene silencing [8]. In gene addition, normal HbA is introduced via lentiviral vectors to SCA patients in order to compensate the reduced oxygen-carriage function of HbS so that new (HbA) and old (HbS) are co-present with near-normal therapeutic status. This mode of gene therapy is realized to be one-time cure of SCA and safer than other methods [9]. Likewise, in gene editing technology the mutated HbS within hematopoietic stem cells is genetically engineered ex vivo and then returned back to the body [10]. Gene silencing uses antisense oligonucleotides complementary to BCL11A mRNA which is responsible for the programming of fetal-to-adult Hb transformation upon adulthood. Hence upon silencing BCL11A, HbF is continued to be expressed resulting in amelioration of SCA complications [11].

A plenty of microRNA (miRNA) were demonstrated to directly and indirectly silence mainstay gene in SCA, BCL11A [12,13]. To our knowledge, targeting erythroid Kruppel-like factor (KLF1), a transcriptional factor regulates erythroid lineage commitment, globin switching, and the end-stage erythrocyte maturation [14], is less-studied. So, the goal of this theoretical work is to find the consensus miRNAs silencing KLF1 gene and design the corresponding miRNA sponge.

## 3 Methodology

### 3.1 Finding consensus KLF1-specific miRNAs

The sequence of KLF1 gene was retrieved from NCBI database having GenBank accession numbers U37106 (gene) and NM_006563 (mRNA). This sequence was searched in 2 databases, namely miRDB [15] (http://www.mirdb.org/) and miRWalk [16] (http://mirwalk.umm.uni-heidelberg.de/) to filter less specific and get the consensus most specific miRNAs. The score threshold was set at 0.95 searching 5’-UTR, CDS and 3’-UTR.

### 3.2 Prediction of secondary structure

The obtained miRNAs secondary structure was predicted using RNAfold web interface [17] (http://rna.tbi.univie.ac.at/cgi-bin/RNAWebSuite/RNAfold.cgi) based on the minimal free energy of folding and partition function. The predicted structures were illustrated in Forna format colored according to the structure type.

### 3.3 Calculation of hybridization energy

The hybridization energy of the found miRNAs and KLF1 3’-UTR (positioned at 100 nucleotides from 3’ end of the gene sequence) was calculated using the online tool RNAhybrid [18] (https://bibiserv.cebitec.uni-bielefeld.de/rnahybrid/). This tool calculates minimum free energy (MFE) of hybridization of miRNA to the best-fitting part of the gene segment (in this case the 3’-UTR). All option parameters were used as default and the MFE values as well as the hybridization representation were tubulated.

### 3.4 Validation of RNAhybrid result

For validation of the RNAhybrid output, RNA-RNA interaction prediction tool IntaRNA 2.0 [19] (http://rna.informatik.uni-freiburg.de/IntaRNA/) was employed. The number of interactions per RNA pair and the minimum number of base-pairs in the seed were adjusted at 5 and 7 respectively.

### 3.5 Design of miRNA sponge

The best obtained miRNA will be elected for design of sponge sequences utilizing miRNAsong server [20] (https://www2.med.muni.cz/histology/miRNAsong/index.php?q=howto) which predicts multiple tandem high affinity binding sites to the desired miRNA. The energy cut-off was set at -25 kcal/mol and using canonical 6-mer seed with 5-nucleotide spacer.

## 4 Results

### 4.1 Finding consensus miRNAs toward KLF1 gene

There are several databases can search miRNA targets and at the same time search miRNA against certain gene. We employed 2 databases (miRWalk and mirDB) so as to get consensus miRNAs toward KLF1 gene. The search process found 6 mutual miRNAs that have silencing activity to KLF1 gene, 3 in the 3’-UTR and 3 in the CDS. 5 out of 6 found miRNAs had a score of 1 indicating the high specificity of the predicted miRNAs to be lead candidate. It is worth noting that the first miRNA (hsa-miR-330-5p) was also confirmed via TargetScan database [21] making it superior candidate.

### 4.2 Secondary structure prediction

The secondary structure of the obtained miRNAs was predicted using RNAfold webserver. Most of them has the regular hairpin-like structure with some bulges at different positions along the structure. The MFE of folding of the first four miRNAs was in the range -1.50 to -5.10 kcal/mol suggesting the high probability of forming acceptable secondary structure except for miRNA 5 and 6 which failed to fold into confined secondary structure (MFE = 0). The predicted secondary structures of all obtained miRNAs are depicted in Fig 2.

**Figure 2.**
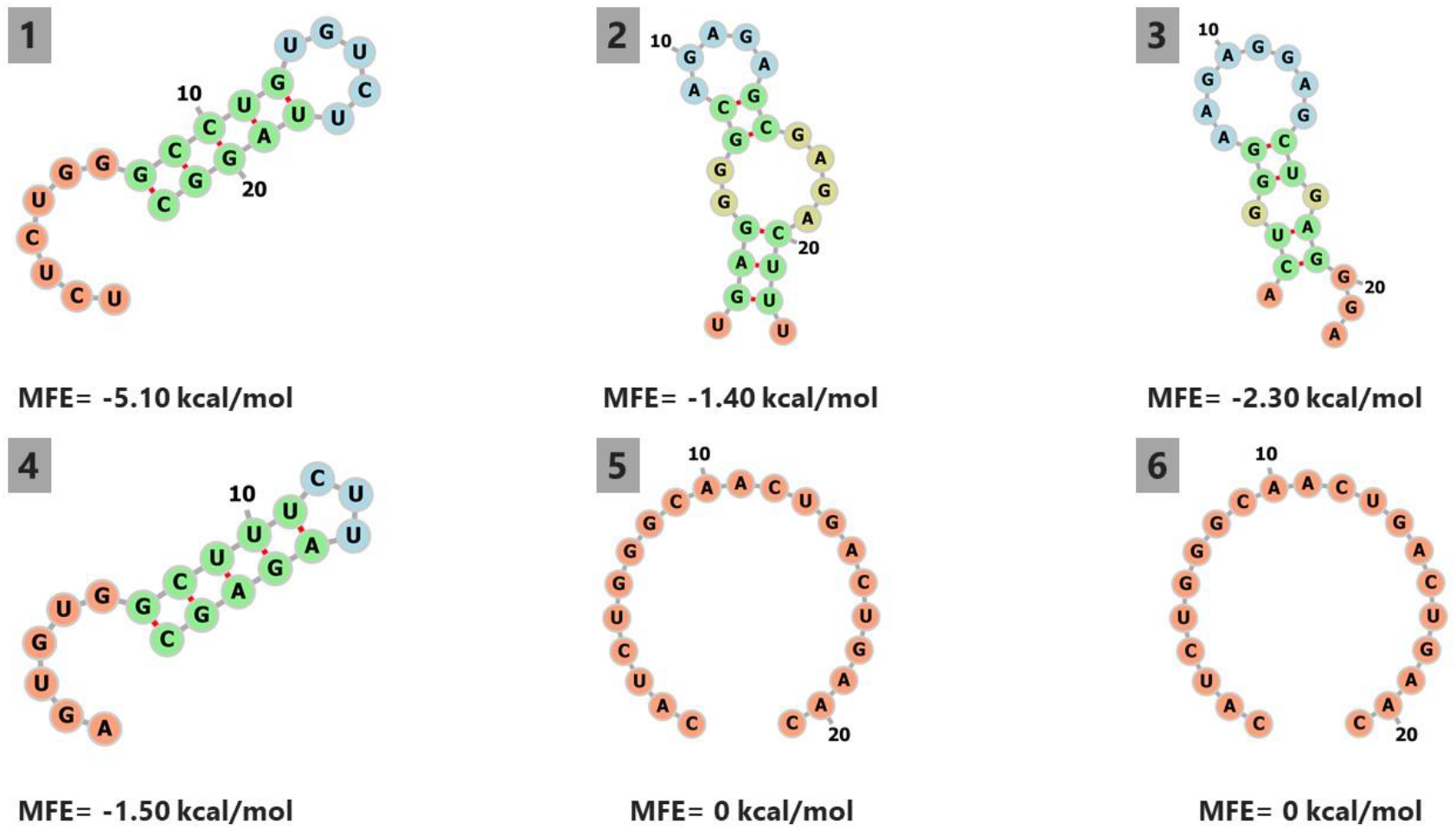
Predicted secondary structures of the filtered miRNAs and their MFE of folding

### 4.3 Hybridization energy

Table 2 elucidates MFE of hybridization between obtained miRNAs and the target gene. The MFE of hybridization ranged between -18.9 (for hsa-miR-644a) to -33.1 kcal/mole (for hsa-miR-330-5p). the miRNAs complementing 3’-UTR was found to have stronger MFE compared to those binds at CDS. The best among all found miRNAs was the first once, hsa-miR-330-5p as suggested by the consensus finding obtained from the 3 searched datasets since the only one approved by TargetScan.

**Table 1.** Obtained consensus miRNAs against KLF1 gne.

| miRNA |  | Score | Binding Site | N Pairings |
| --- | --- | --- | --- | --- |
| **3'-UTR** | hsa-miR-330-5p | 1 | 166,187 | 18 |
|  | hsa-miR-423-5p | 1 | 11,031,125 | 17 |
|  | hsa-miR-4646-5p | 1 | 85,126 | 17 |
| **CDS** | hsa-miR-644a | 1 | 13,741,387 | 12 |
|  | hsa-miR-1298-3p | 1 | 14,421,473 | 16 |
|  | hsa-miR-6729-3p | 0.98 | 14,621,504 | 16 |

3'-UTR: 3'-untranslated region, CDS: coding sequence

**Table 2.**
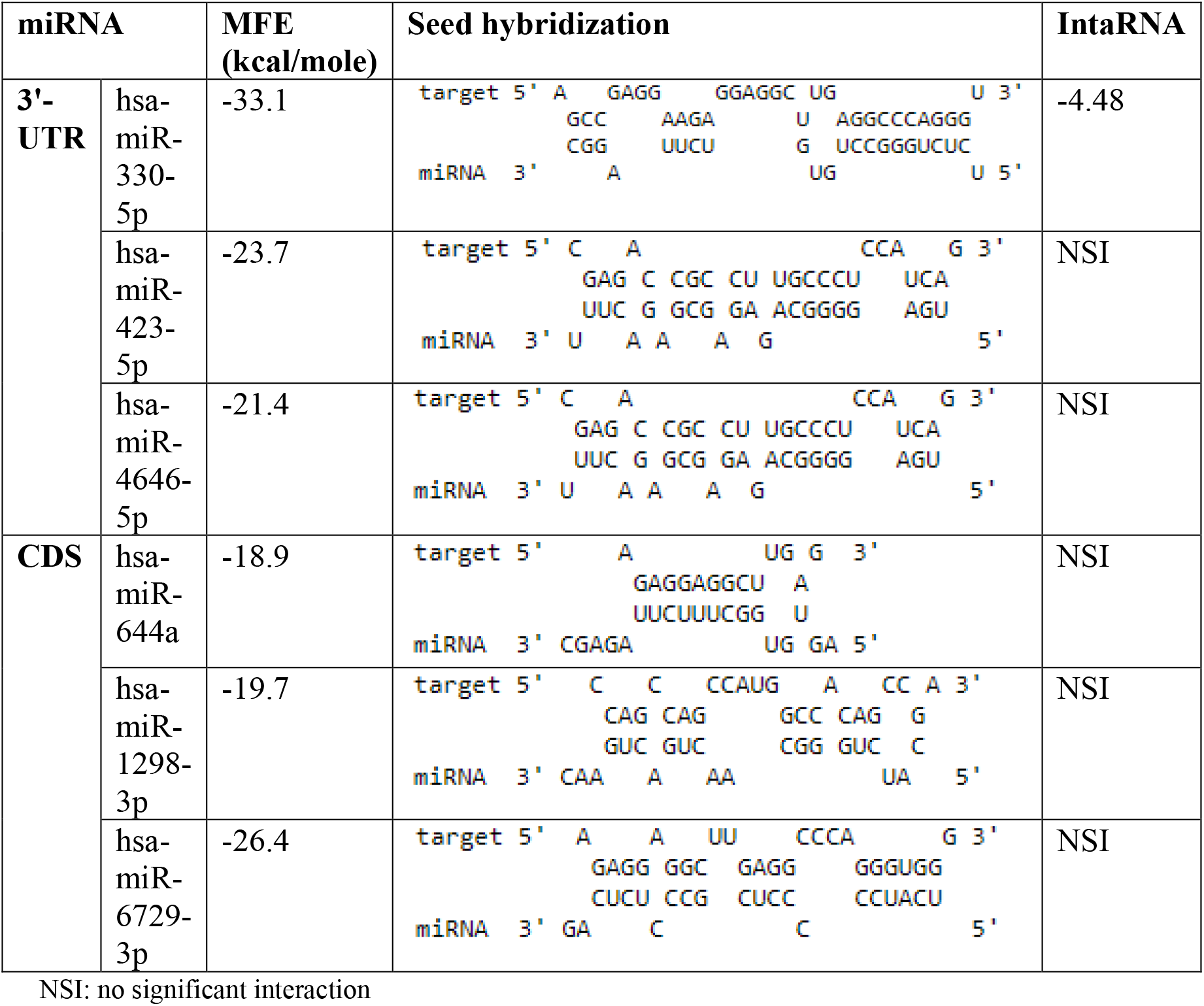
Calculated hybrizidation energies of miRNAs and KLF1 3’-UTR

In order to validate the hybridization data obtained using RNAhybrid between the miRNA sequences and KLF1 transcript, RNA-RNA interaction prediction tool IntaRNA 2.0 was employed. Interestingly, only the first miRNA, hsa-miR-330-5p, gave significant interaction thus validating the RNAhybrid results which ranked it as the top 1 in terms of hybridization energy. KLF1-hsa-miR-330-5p interaction yielded a hybridization energy -20.04 kcal/mol but the folding energy of the KLF1 (10.45) and miRNA (5.11) have lowered the total hybridization energy to -4.48 kcal/mol.

### 4.4 Prediction of miRNA sponge

By applying the 5-nucleotide spacer (UGUGU) and 9-13 nucleotide bulge, the predicted hsa-miR-330-5p sponge gave a total free energy of duplex -72.8 kcal/mol. The miRNA sponge generated 2 high affinity binding sites with free energy -36.7 and -36.1 kcal/mole, respectively (Fig 3).

**Figure 3.**
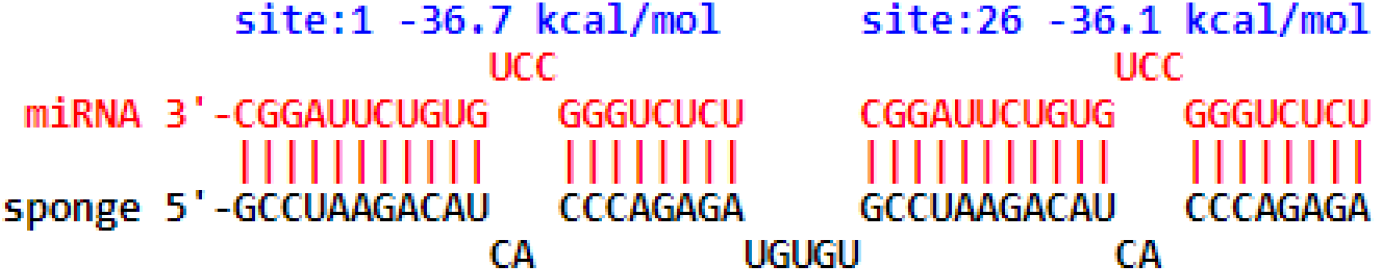
Designed hsa-miR-330-5p sponge showing hybridization pattern and energy

Overall, hsa-miR-330-5p proved itself as the best candidate miRNA to be a lead target to be hybridized and thus inhibited by the sponge sequence (GCCTAAGACATCACCCAGAGATGTGTGCCTAAGACATCACCCAGAGA) for a variety of reasons: (i) yielded the strongest hybridization energy in RNAhybrid, (ii) the only miRNA which yielded significant hybridization using IntaRNA 2.0, and (iii) the only miRNA detected by TargetScan dataset.

## 5 Discussion

SCA is one of the most severe inherited diseases that thought to affects 3-6 million people in the world, 70% of whom lives in African continent due to limited health care accessibility [22]. Many of children born with this disease die sooner compared to healthy ones because of the devastating consequences of the disease. This includes stroke and renal failure and hyposplensim-opportunistic infections [23]. miRNA have been shown to do controlling actions in erythrocytes, platelets, and leukocytes. Significantly Differentially expressed miRNAs were found in sickled Hb (HbS) including miRNA-299, miRNA-144, miRNA-140, miRNA-451 whereas miRNA-320, let-7s, miRNA-181, miRNA-141 were detected in normal mature Hb (HbA) [24]. Dysregulation of certain miRNAs have been reported to pose ameliorating effects in some circumstances and increases the severity of SCA in other situations. miRNAs perform pivotal roles within erythrocytes such as regulation of cell cycle, iron concentrations, oxidative stress, hemolysis process and, most notably, triggers HbF or HbA expression and transformation of the genotype expressed [13]. One of the switching transcriptional factors is KLF1. In the normal state, KLF1 upregulates β-globin expression in the developing erythrocyte precursors during adulthood contributing to the emergence and progression of HbA (which is mutated in SCA) in place of HbF (normally functioning even in HbS) [25]. Gillinder et al. [26] used gene editing technology via CRISPR-Cas9 to knockout KLF1 gene as a means to cure SCA in human cells *in vitro*. γ-globin was upregulated 10-fold, BCL11A down-regulated 3-fold, and HbF levels were detected to be between 40-60% of total Hb. Moreover, KLF1 ablation was accompanied by downregulation of ICAM-4 and BCAM, cellular adhesion molecules implicated in triggering vaso-occlusive episodes. The results of the current work candidates hsa-miR-330-5p as a superior miRNA that predicted to control KLF1 translation. Therefore, generating miRNA sponge (GCCTAAGACATCACCCAGAGATGTGTGCCTAAGACATCACCCAGAGA) to this miRNA could achieve similar findings obtained from Gillinder et al. [26] given that gene silencing is safer and readily applicable than gene editing [27].

## 6 Conclusion

We have searched 2 miRNA databases (miRDB and miRWalk) to find the consensus miRNAs targeting KLF1 gene and validated the results by third database (TargetScan). Together with hybridization tools, hsa-miR-330-5p was predicted to be the most common and implicated miRNA toward KLF1 translation as a means to genetically treat SCA by utilizing the corresponding miRNA sponge. So, it is suggested to examine the efficacy of this in silico finding via wet-lab setting.

